# RNA *in situ* conformation sequencing reveals novel long-range RNA structures with impact on splicing

**DOI:** 10.1101/2022.11.07.515435

**Authors:** Sergei Margasyuk, Marina Kalinina, Marina Petrova, Dmitry Skvortsov, Changchang Cao, Dmitri D. Pervouchine

**Affiliations:** Skolkovo Institute of Science and Technology, Moscow 143026, Russia; Moscow State University, Faculty of Chemistry, Moscow 119991, Russia; Key Laboratory of RNA Biology, Institute of Biophysics, Chinese Academy of Sciences, Beijing 100101, China

**Keywords:** long-range, RNA interaction, RIC-seq, PCCR, PHF20L1, CASK, splicing

## Abstract

Over past years, long-range RNA structure has emerged as a factor that is fundamental to alternative splicing regulation. Since an increasing number of human disorders are now being associated with splicing defects, it is essential to develop methods that assess long-range RNA structure experimentally. RNA *in situ* conformation sequencing (RIC-seq) is the method that recapitulates RNA structure within physiological RNA-protein complexes. In this work, we juxtapose RIC-seq experiments conducted in eight human cell lines with pairs of conserved complementary regions (PCCRs) that were predicted *in silico*. We show statistically that RIC-seq support strongly correlates with PCCR properties such as equilibrium free energy, presence of compensatory substitutions, and occurrence of A-to-I RNA editing sites and forked eCLIP peaks. Based on these findings, we prioritize PCCRs according to their RIC-seq support and show experimentally using antisense nucleotides and minigene mutagenesis that PCCRs in two disease-associated genes, *PHF20L1* and *CASK*, impact alternative splicing. In sum, we demonstrate how RIC-seq experiments can be used to discover functional long-range RNA structures, and particularly those that regulate alternative splicing.

## Introduction

Double-stranded RNA structures, many of which are long-range and span thousands of nucleotides, provide an additional layer in the pre-mRNA splicing regulation (*1*). Understanding of mechanisms that underlie pre-mRNA splicing is not only an outstanding fundamental question of molecular biology but also a key element for the development of effective therapies in splicing-associated human diseases (*2–4*). Yet, systematic experimental assessment of long-range RNA structures has not been possible until recent development of high-throughput sequencing protocols based on RNA proximity ligation (*5*).

RNA-RNA interactions can be assessed *in vivo* and *in vitro* by a number of strategies, which can be broadly subdivided into classical RNA structure probing and conformation capture type of methods (*6, 7*). The use of classical RNA structure probing for long-range RNA structures is limited because it can only tell whether an RNA base is paired, but it cannot tell to which other base. In contrast, conformation capture methods employ digestion and religation of RNAs crosslinked in macromolecular complexes to assess spatial proximity of RNA strands and, therefore, are able to identify base-paired regions. For instance, PARIS (*8*), LIGR-seq (*9*), SPLASH (*10*), and COMRADES (*11*) specifically target double-stranded regions using psoralen derivatives, which intercalate into RNA duplexes and induce reversible crosslinking. However, a new class of RNA *in situ* conformation sequencing (RIC-seq) methods, in which RNAs are crosslinked through RNA-binding proteins (RBPs), are increasingly being employed to identify RNA structures that occur within physiological RNA-RBP complexes (*12, 13*).

While computational prediction of long-range RNA structures at the scale of full eukaryotic transcripts is problematic, it still can be approached by comparative methods that measure the rate of compensatory substitutions in evolutionarily conserved regions (*14*). Our recent study cataloged 916,360 pairs of conserved complementary regions (PCCRs) in human protein-coding genes that have a remarkable association with splicing and strong support by RNA editing sites and the so-called forked eCLIP peaks, i.e., simultaneous RBP footprints near complementary sequences (*15*). However, only a fraction of structural alignments are convincingly supported by covariations, while the majority of PCCRs including RNA structures with established biological function lack evolutionary support because the amount of sequence variation is insufficient to estimate statistical significance through compensatory substitutions (*15*). This naturally leads to a question of whether computational RNA structure prediction can be improved by taking into account RNA proximity ligation data.

In this work, we study the problem of assessment and prioritization of PCCRs using a panel of RIC-seq experiments. We show that local RIC-seq support strongly correlates with important PCCR properties such as equilibrium free energy, significant compensatory substitutions, the presence of A-to-I RNA editing sites, and forked eCLIP peaks. Based on RIC-seq evidence, we select two disease-associated human genes and demonstrate experimentally that PCCRs strongly impact their alternative splicing. One of these genes, the plant homeodomain finger protein 20-like 1 (*PHF20L1*) encodes a histone methylation reader that interacts with mono- and dimethylated lysines in H3K4me1, H4K20me1, H3K27me2, as well as with epigenetic factors, e.g., *DNMT1* (*16–18*). It is also involved in maintaining the stability of methylated *SOX2* and *pRb* proteins (*19, 20*). *PHF20L1* is essential for epigenetic inheritance in mammals, cell pluripotency and differentiation, and the maintenance of the G1-S phase checkpoint, while aberrations of its expression are common for breast, colorectal, and ovarian cancers (*21–23*). The other gene, *CASK*, encodes a human calcium/calmodulin-dependent serine protein kinase, however the protein does not possess kinase activity (*24*) and functions as a scaffolding protein involved in presynaptic and postsynaptic transmission (*25, 26*). In mice, the deletion of *CASK* is lethal (*27*), while its inactivation in pancreatic *β* cells affects glucose homeostasis and insulin sensitivity (*28*). *CASK* interacts with *Tbr-1*, a T-box transcription factor involved in forebrain development (*29*) and possibly playing a role in epithelial cell polarity establishment in mammals (*30*).

Our experimental strategy was to probe a PCCR using antisense oligonucleotides (AONs) obstructing one or the other complementary sequence, and then use site-directed mutagenesis in minigenes to prove that long-range RNA structure indeed impacts alternative splicing (*31*).

## Results

### Concordance of PCCRs and RNA contacts

To evaluate the support of PCCRs by RNA contacts, we analyzed a panel of RIC-seq experiments conducted in eight human cell lines including GM12878, H1, HeLa, HepG2, IMR90, K562, hNPC, and *in vitro* differentiated neuronal cells (see Methods). We hypothesized that digestion and religation of RNA strands adjacent to stretches of complementary nucleotides should result in RNA contacts near PCCRs as shown in Figure 1A, i.e., supporting them from the outer or from the inner side, which correspond to split reads in the collinear (2-3) or chimeric (1-4) orientation, respectively.

**Figure 1:**
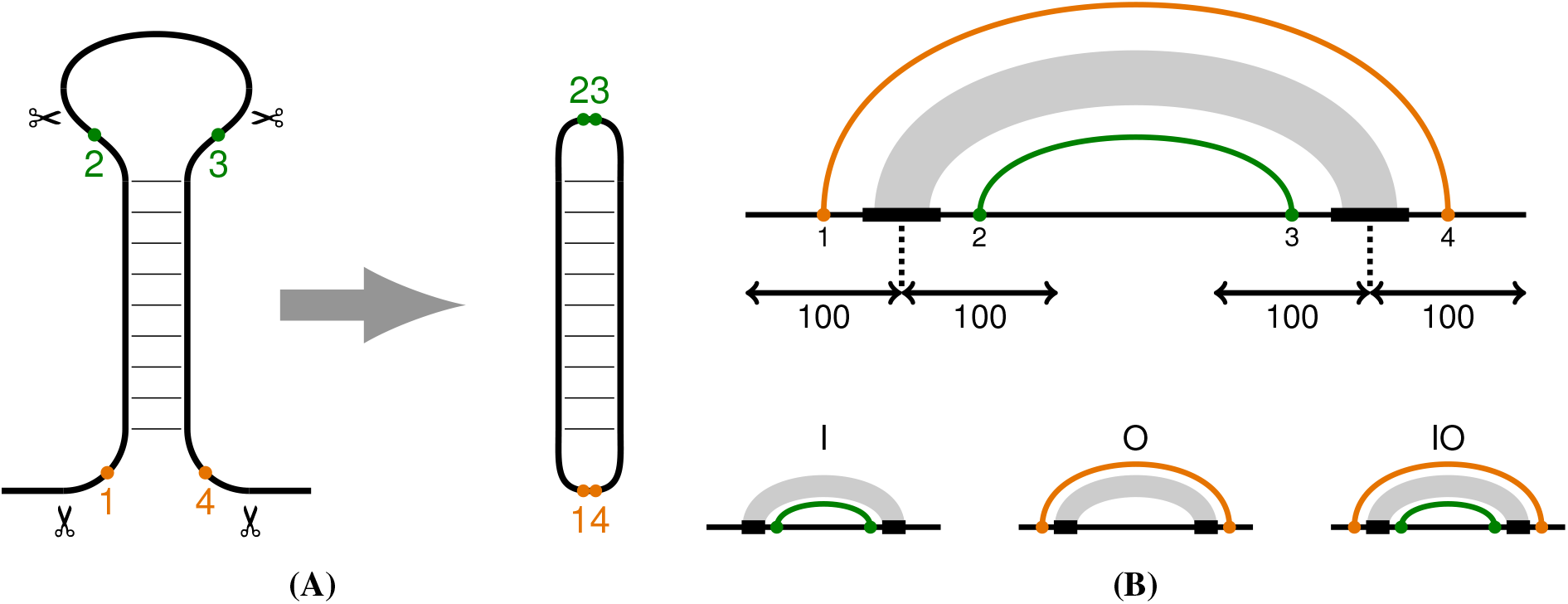
RNA contacts. **(A)** Digestion and religation of RNA strands adjacent to a PCCR result in RNA contacts (in the form of split reads) supporting it from the inner (2-3) or the outer (1-4) side. **(B)** The inner (I) and the outer (O) contacts correspond to the inner and outer arcs with respect to the RNA structure. The inner contacts are represented by chimeric split reads, as in circular RNAs.

We selected 802,536 out of 916,360 PCCRs documented in (*15*) with the distance between complementary regions of at least 200 nts and surrounded them by windows with the radius 100-nt (Figure 1B). As a result, PCCRs were subdivided into four mutually exclusive groups: ones supported by at least one RNA contact in a 100-nt window inside but not outside (I), ones supported by at least one RNA contact outside but not inside (O), ones supported by at least one RNA contact both inside and outside (IO), and the rest of the PCCRs (N). At this point we chose to analyze PCCR support only using presence/absence call, i.e., regardless of the number of supporting RIC-seq reads because each individual RIC-seq experiment yields sparse RNA contacts (*32*). The subdivision into IO, I, O, and N groups was done at the level of each individual RIC-seq experiment, as well as for the union of bioreplicates in each cell line, and also for the pooled set of all RIC-seq experiments in all cell lines (Table 1). Remarkably, the number of PCCRs in the pooled IO group was much larger than expected additive contribution from individual cell lines, i.e., many PCCRs are supported from inside in one cell line and from outside in another. This indicates that a collective evidence from multiple datasets may be more appropriate than that limited to a particular biological condition.

**Table 1:** The number of PCCRs in the IO, I, O, and N groups (see text). “Pooled” represents PCCR groups with respect to the pooled set of contacts from all cell lines.

| Cell | IO | I | O | N |
| --- | --- | --- | --- | --- |
| GM12878 | 3,099 | 13,333 | 8,326 | 777,778 |
| H1 | 1,614 | 10,058 | 6,129 | 784,735 |
| HeLa | 673 | 4,773 | 2,864 | 794,226 |
| HepG2 | 2,061 | 9,169 | 6,420 | 784,886 |
| IMR90 | 1,455 | 7,842 | 4,523 | 788,716 |
| K562 | 2,390 | 11,127 | 7,920 | 781,099 |
| hNPC | 960 | 6,504 | 3,929 | 791,143 |
| Neuron | 3,080 | 13,941 | 10,225 | 775,290 |
| Pooled | 19,852 | 37,505 | 24,284 | 720,895 |

To estimate the concordance between bioreplicates, we first estimated the overlap between PCCRs groups in two bioreplicates of each RIC-seq experiment. As expected, the largest overlap was observed for the cognate classes in the IO, I, and O groups (Figure S1). Next, we compared IO, I, and O classes between RIC-seq replicates in different cell lines. The groups were most similar between bioreplicates, but the degree of similarity was expectedly lower when comparing cell lines with each other (Figure S2). The level of consistency 9–15% between bioreplicates indicated once again that each individual RIC-seq experiment yields sparse data, and that aggregating results from several independent RIC-seq experiments could provide a better strategy for further analysis.

### Properties of PCCRs supported by RNA contacts

PCCRs are characterized by a number of factors including the equilibrium free energy (Δ*G*), which is estimated from the parameters of thermodynamic RNA folding (*33*), the *E*-value, which scores independent compensatory substitutions on a phylogenetic tree (*34*), the overlap with A-to-I RNA editing sites, which are indicative of double-stranded RNA context (*35, 36*), and the occurrence of forked eCLIP peaks (*15*), which reflect RBP crosslinking near complementary RNA strands (Figure S3).

To evaluate the relationship between these factors and the observed RNA contacts, we first identified PCCRs that are supported by at least one RNA contact in the IO, I, and O groups in at least *k* cell lines and compared them to PCCRs that were not supported by RNA contacts (i.e., *k* = 0). The stability of PCCRs (i.e., the absolute value of Δ*G*), the proportion of PCCRs with A-to-I RNA editing sites, and the proportion of PCCRs with forked eCLIP peaks consistently increased with increasing *k*, while the compensatory substitution support (*E*-value) did not show a strong dependence on *k* (Figures 2, S4 and S5). To further characterize PCCRs with different levels of RIC-seq support, we estimated the number of PCCRs that are supported by I and O contacts using a bivariate number of cell lines and constructed 95% confidence intervals for their equilibrium free energy (Tables S1 and S2). We observed a consistent trend of increasing Δ*G* (by absolute value) with increasing RIC-seq support from either side, with sufficiently many PCCRs having very strong support, e.g., 534 PCCRs being supported in at least five cell lines by both inside and outside contacts.

**Figure 2:**
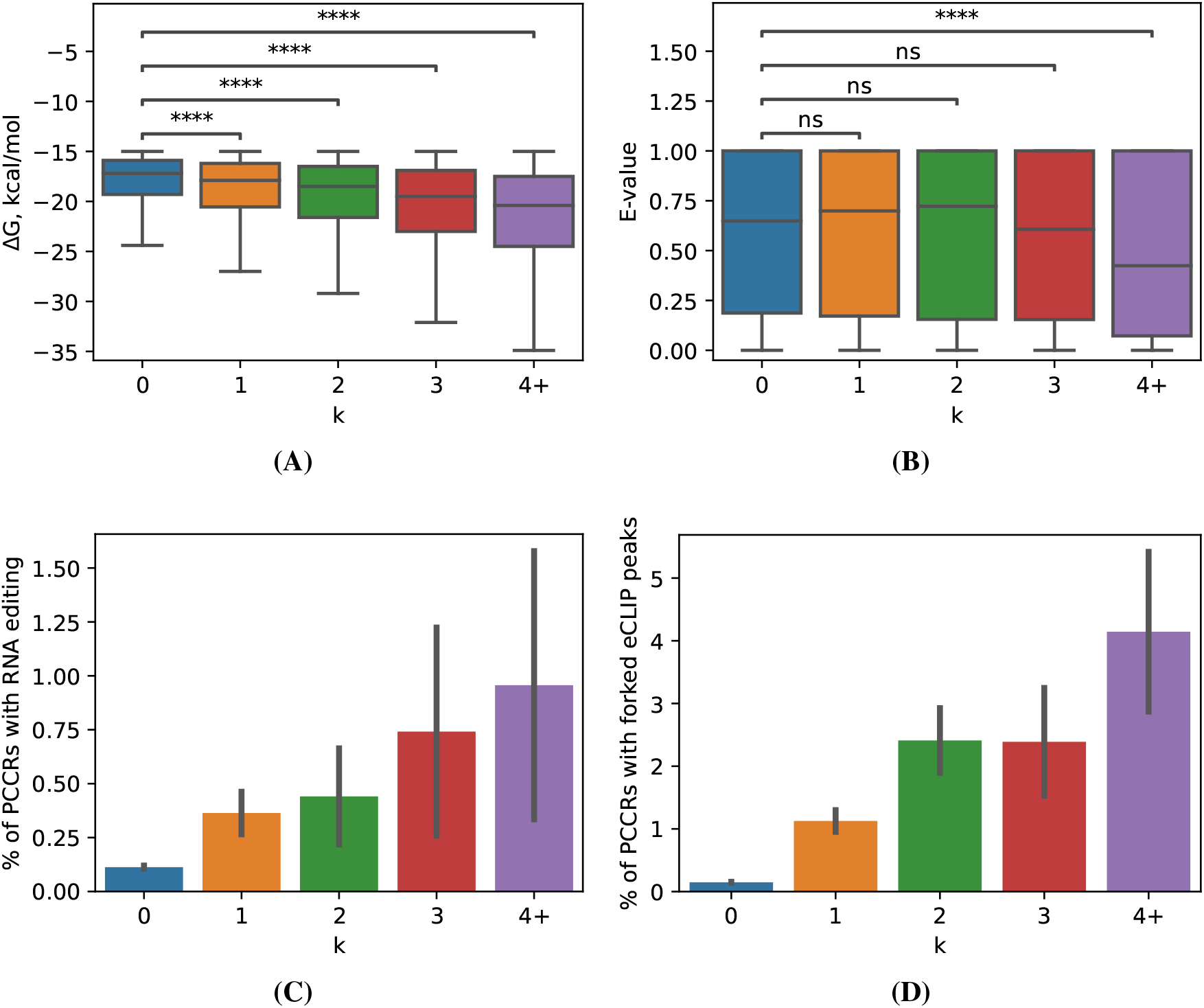
PCCR properties. The equilibrium free energy Δ*G* (A), the R-scape *E*-value (*34*) (B), the frequency of A-to-I RNA editing sites (C), and the frequency of forked eCLIP peaks near PCCRs supported from inside and outside (IO category) in at least *k* cell lines. Statistically discernible differences at the 5%, 1%, 0.1%, and 0.01% significance level are denoted by ^*^, ^**^, ^***^, and ^****^, respectively (two-tailed Mann-Whitney test). Non-significant differences are denoted by ‘ns’. Whiskers indicate 95% confidence intervals for proportions.

Next, we focused on exons located inside PCCRs and estimated their median inclusion rates (percent spliced in, PSI, or Ψ) in RNA-seq experiments conducted in a matched set of cell lines. As expected, the distribution of Ψ consistently shifted towards lower values with increasing *k* (Figure 3A) in agreement with previous findings that exon inclusion drops with increasing stability of the surrounding PCCR (*15*). To dissect differential alternative splicing, we considered PCCRs that were supported by at least four reads in at least three cell lines and compared the inclusion rate of exons looped out by these PCCRs in the experiments with and without RIC-seq support. The difference of exon inclusion rates in cell lines with vs. without RIC-seq support (ΔΨ) was significantly shifted towards negative values (Wilcoxon test, P-value = 10^−9^) indicating that alternative splicing of an exon can be modulated by PCCRs assembling and disassembling in different conditions (Figure 3B). In the interest of future experimental studies, PCCRs with differential RIC-seq support are listed in Supplementary Data File 1 along with exon inclusion rates.

**Figure 3:**
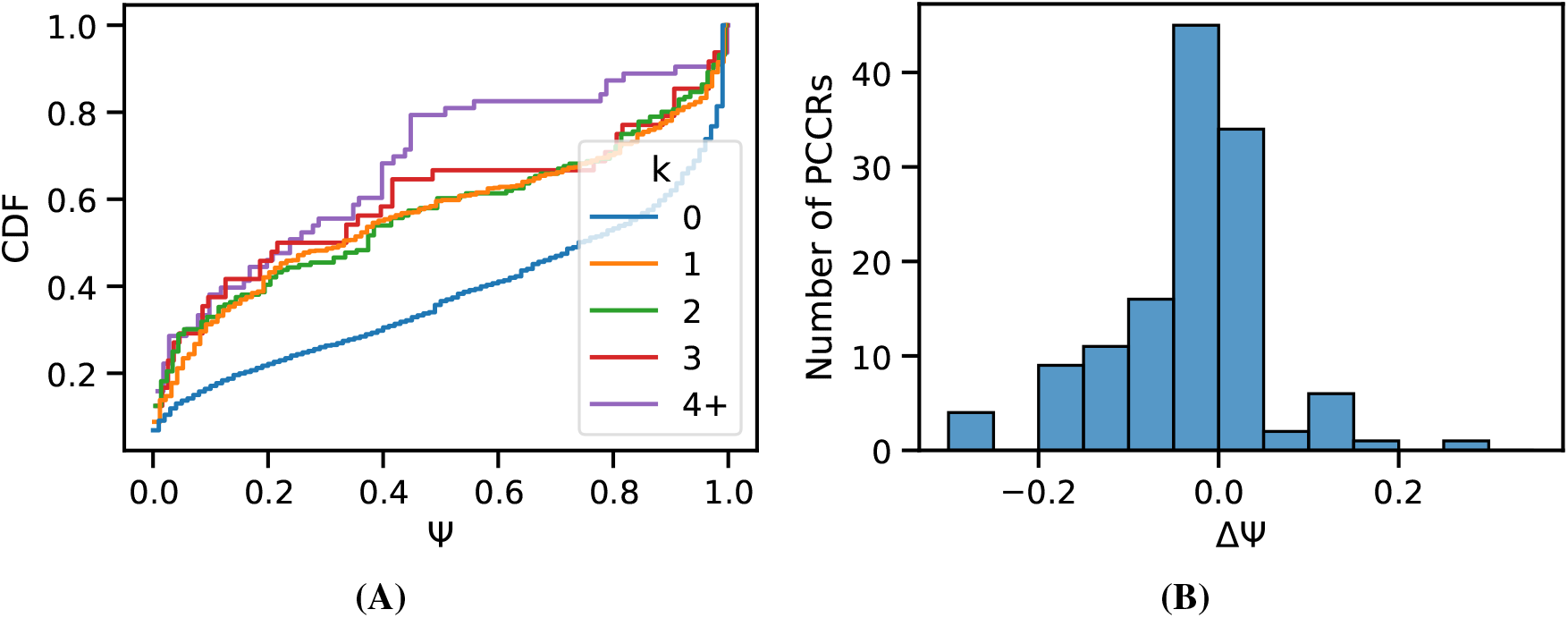
Exons inside PCCRs. **(A)** The distribution of average exon inclusion rate (Ψ) for exons in PCCRs supported both inside and outside (IO category) in at least *k* cell lines. CDF represent cumulative probability density function. **(B)** The distribution of ΔΨ = Ψ_*h*_ − Ψ_*l*_, where Ψ_*h*_ and Ψ_*l*_ are the average Ψ values in cell lines with and without RIC-seq support, respectively.

To further characterize the relationship between RNA contacts and PCCRs, we constructed a random forest (RF) classifier (see Methods) to predict the occurrence of forked eCLIP peaks nearby. Towards this goal, we surrounded each complementary region, left (*L*) and right (*R*), by three 50-nt bins extending upstream (−3, −2, −1) and downstream (1, 2, 3) from the center of the region (Figure 4A). Since the density of RNA contacts is higher for shorter contacts, we also included the distance between the complementary parts, also referred to as PCCR spread, as a variable in the model (*32*). PCCR spread alone was able to predict eCLIP peaks near PCCR with *AUC* = 0.66, while the addition of RNA contacts increased the quality of the predictions to *AUC* = 0.74 (Figure 4B). The presence of immediately adjacent inner and outer contacts, i.e., contacts between bins 1*L* and −1*R*, and contacts between bins −1*L* and 1*R*, were the two most important features for the classifier (Figure 4C). This effect was not due to higher density of RNA contacts at shorter distances because PCCR spread was included as a covariate in this analysis.

**Figure 4:**
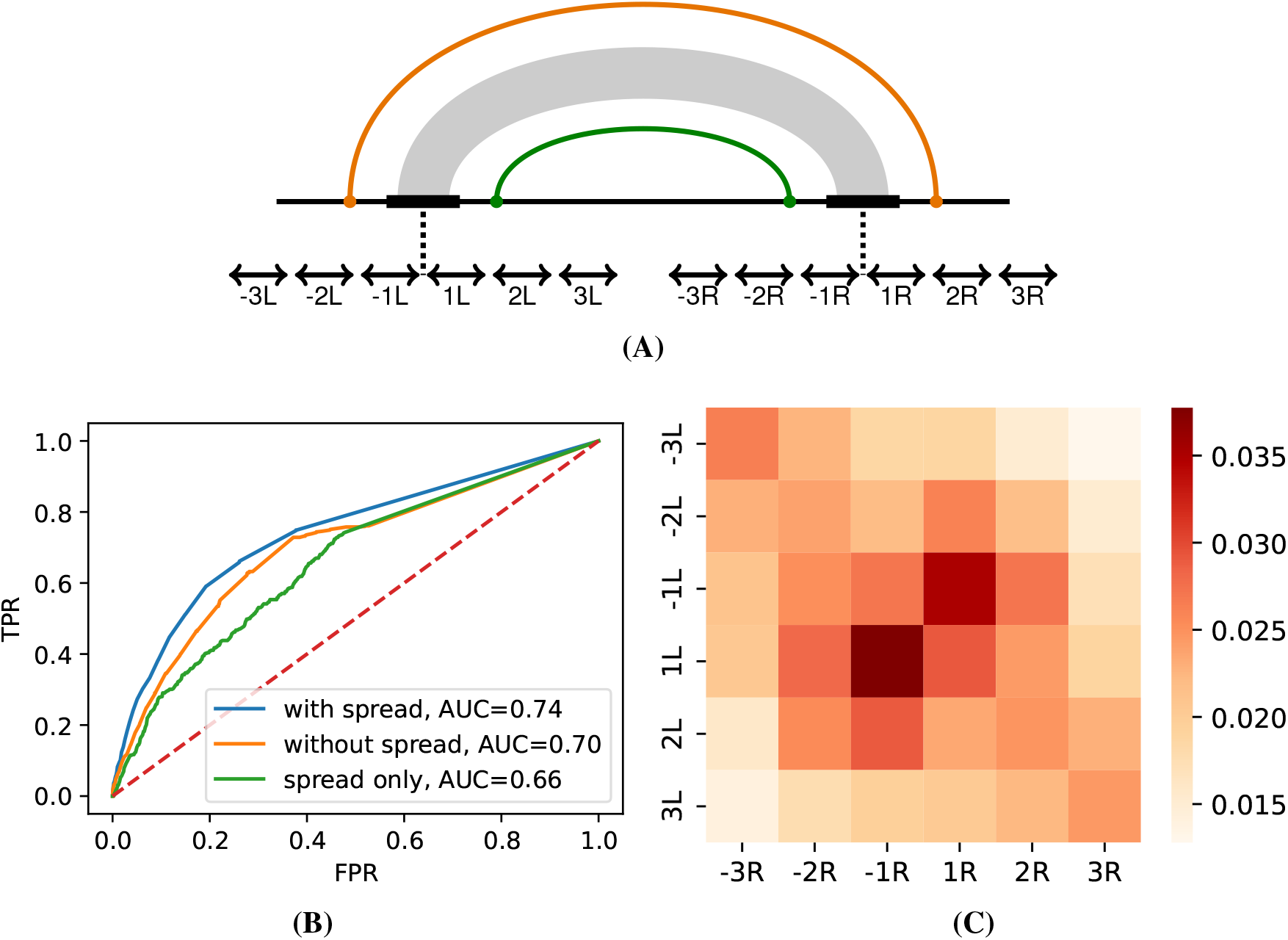
Random forest (RF) classifier. **(A)** Complementary sequences of each PCCR were surrounded by six 50-nt bins centering in the middle of the despaired region, and split read counts representing RNA contacts from RIC-seq were computed for each of the 36 combinations. **(B)** The performance (TPR, true positive rate, vs. FPR, false positive rate) of the RF classifier predicting the presence of forked eCLIP peaks as a function of (1) PCCR spread, (2) split read counts, and (3) both spread and split read counts. AUC is the area under the curve. **(C)** Feature importance for the 36 combinations of bins. The two most important features are RIC-seq contacts between 1*L* and −1*R* and between −1*L* and 1*R* bins, which correspond to the inner and outer contacts immediately adjacent to PCCR.

Taken together, these observations suggest that RIC-seq support is correlated with important PCCR properties, and that correlation is localized to the regions adjacent to PCCRs. Thus, we created a table listing 115,162 PCCRs supported by RIC-seq in at least one cell line (Supplementary Data File 2). From this list, we selected two candidates, a PCCR with Δ*G* = −23.6 kcal/mol in the *PHF20L1* gene (id838701) and a PCCR with Δ*G* = −21.6 kcal/mol in the *CASK* gene (id902118), on the basis of RIC-seq support (in at least two cell lines both inside and outside), evolutionary conservation scores (*s*_3_), presence of a cassette exon inside PCCR, feasibility of cloning and expression, gene expression level, exon inclusion rate, and other considerations (*15*). In the next section, we present experimental validation of the impact of these PCCRs on splicing using antisense oligonucleotides and site-directed mutagenesis on minigenes in human and murine cell lines.

## Case studies

### PHF20L1

The *PHF20L1* gene produces two transcript isoforms, *PHF20L1*-a and -b, which differ by the inclusion of alternative cassette exon 6 (*18*). Introns flanking exon 6 contain a pair of conserved complementary regions, R1 and R2, which could form stable RNA structure (Δ*G* = −23.6 kcal/mol) and contribute to the regulation exon 6 alternative splicing (Figure 5A). This PCCR was identified as supported by RNA contacts observed in RIC-seq experiments in neuronal, HeLa, and HepG2 cell lines (Figure S7A).

**Figure 5:**
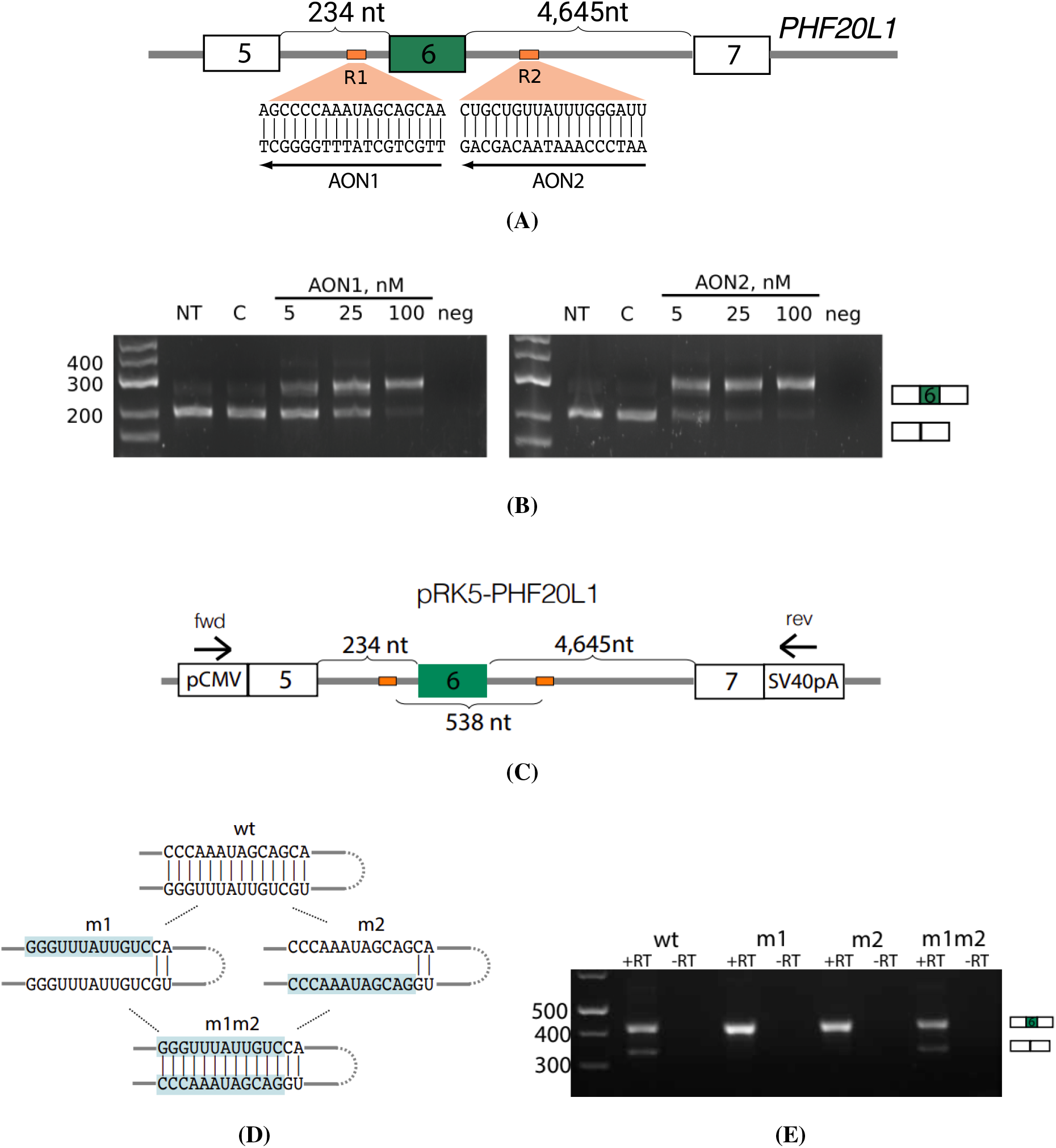
PHF20L1. **(A)** Cassette exon 6, complementary sequences R1 and R2, and their respective AONs (AON1 and AON2). **(B)** The addition of 5 nM of AON1 or AON2 is sufficient to suppress exon 6 skipping. **(C)** The scheme of a minigene expressing a fragment of the *PHF20L1* gene. Primer locations are indicated by arrows. **(D)** Mutagenesis in the minigene. In m1 and m2 mutants, the base pairing between R1 and R2 is destroyed, while in the compensatory double mutant m1m2, it is restored. **(E)** Exon 6 skipping in m1 and m2 mutants is suppressed, but in m1m2 it is reverted to that of the WT (See also quantitative analysis in S7).

To evaluate the role of R1/R2 base pairing in alternative splicing of the endogenous transcripts, we designed AONs complementary to R1 and R2, called AON1 and AON2, respectively (Figure 5A) and measured the rate of exon 6 inclusion in response to increasing AONs concentrations. Both qualitative and quantitative RT-PCR analysis (Figures 5B S7B, and S7C) confirmed that 5 nM or higher concentration of either of two AONs was sufficient to substantially increase exon 6 usage in the endogenous transcript.

Next, we constructed a minigene that contains a part of *PHF20L1* gene spanning from exon 5 to exon 7 (Figure 5C) and introduced mutations that disrupted and restored base pairings between R1 and R2 (Figure 5D). Mutations disrupting R1/R2 base pairing, called m1 and m2, promoted exon 6 inclusion, while the compensatory double mutant m1m2, which restores R1/R2 base pairing, also reverts the splicing pattern to that of the wild type (WT) both qualitatively (Figure 5E) and quantitatively (Figure S7D). The response of exon 6 inclusion to AONs and reversal of the WT splicing in the compensatory double mutant strongly indicate that exon 6 inclusion is controlled by R1/R2 base pairing.

### CASK

Alternative splice isoforms of *CASK* differ by the inclusion or skipping of cassette exon 19, which was suggested to modulate *CASK* binding to other proteins across developmental stages and in cell populations with different neuronal activity (*37,38*). Introns flanking exon 19 contain R3 and R4, a PCCR located more than 3,000 nts apart from each other and potentially looping out exon 19 (Figure 6A). The R3/R4 interaction is supported by RIC-seq contacts in IMR90, hNPC, neuronal, and GM12878 cell lines (Figure S8A).

**Figure 6:**
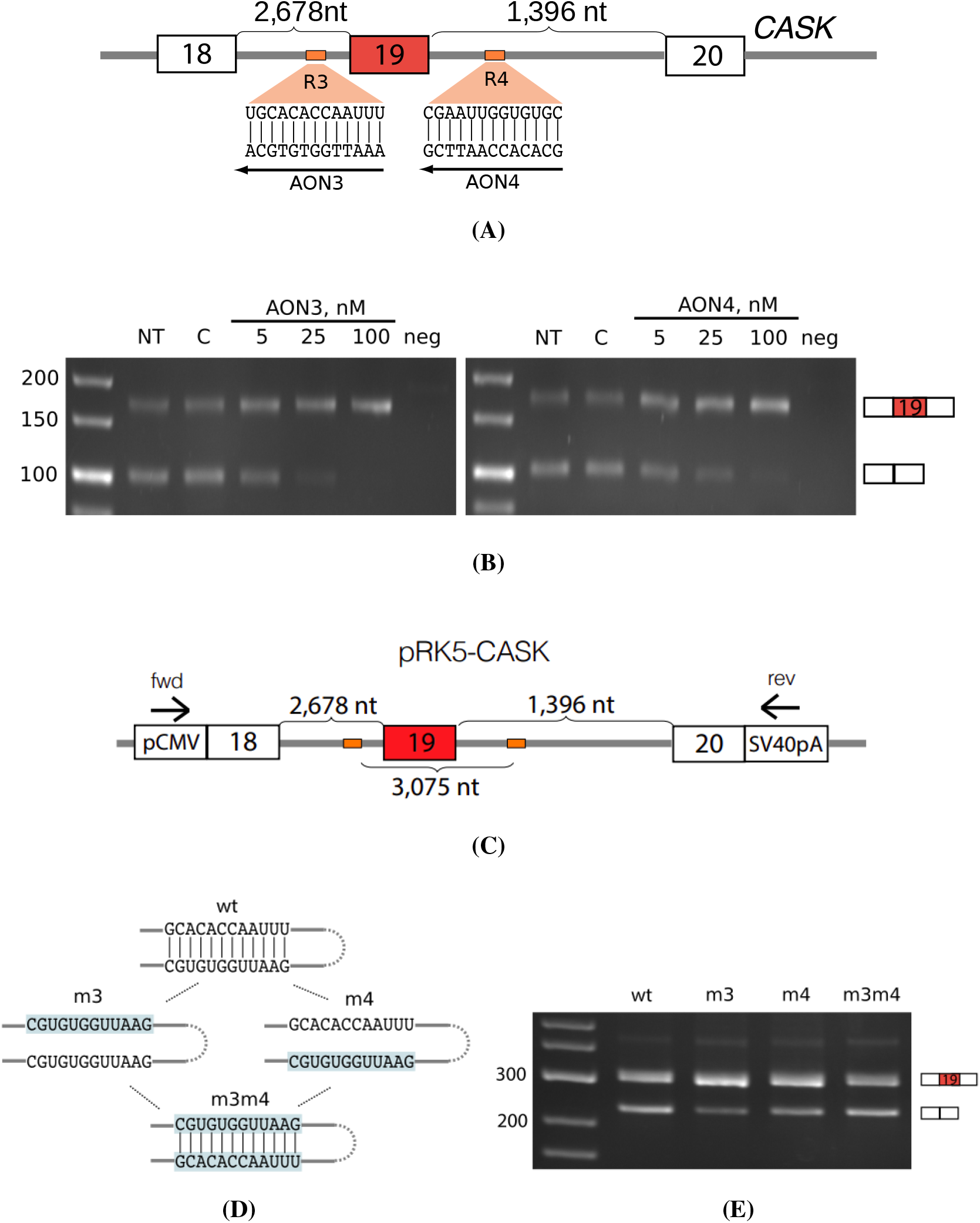
CASK. **(A)** Cassette exon 19, complementary sequences R3 and R4, and their respective AONs (AON3 and AON4). **(B)** Exon 6 skipping is almost completely abolished with the addition of 100 nM of AON1 or AON2. **(C)**, **(D)**, and **(E)** are as in Figure 5. See also quantitative analysis in Figure S8 and mouse *CASK* in Figure S9 and S10.

Exon 19 inclusion rate substantially increased in response to the treatment by AON3 and AON4, LNA complementary to R3 and R4, respectively (Figures 6B, S8B, and S8C). Next, we assembled a minigene that contains a fragment of *CASK* spanning between exons 18 and 20 (Figure 6C) and introduced disruptive and compensatory mutations as before (Figure 6D). Single mutations (m3 and m4) expectedly led to the loss of exon skipping, while the compensatory mutant (m3m4) restored the exon inclusion rate of the WT (Figure 6E, Figure S8D).

Since R3 and R4 are highly evolutionarily conserved, we next chose to interrogate murine *CASK* gene using the same AONs. The rate of exon 19 inclusion in NIH 3T3 mouse fibroblasts increased by on average 40% with increasing AON3 and AON4 concentration (Figure S9). Furthermore, we constructed a minigene carrying a fragment of murine *CASK* gene and applied the same mutagenesis strategy (Figure S10A and Figure 6D). Again, single mutants (m3 and m4) promoted exon 19 inclusion, while compensatory double mutant (m3m4) recovered the WT splicing ratio, however not completely (Figure S10B and S10C). Taken together, these results indicate that complementary base pairings between R3 and R4 in the human and murine *CASK* genes represent an evolutionarily and functionally conserved mechanism controlling exon 19 alternative splicing.

## Discussion

The structure of eukaryotic transcripts has a remarkably complex multi-level organization, in which local secondary structure elements are hierarchically assembled into a tertiary structure that is stabilized by RBPs and long-range intramolecular base pairings (*14*). RIC-seq technology has made it possible to take snapshots of this complex and dynamic picture by detecting RNA-RNA contacts at single-base resolution (*12, 13*). These contacts, however, do not necessarily correspond to RNA structure but rather constitute the fact that two RNA strands are located proximally in 3D, possibly due to intra- and intermolecular RNA-RNA interactions, or possibly due to interaction mediated by proteins. For instance, *hnRNPA1* and *PTB* splicing factors bound to the pre-mRNA tend to form dimers, thus creating RNA contacts that are not mediated by base pairings (*39, 40*).

The analysis presented here interrogates the relationship between intramolecular RNA contacts observed via RIC-seq in a panel of cell lines and PCCRs obtained through bioinformatic predictions, thus likely capturing RNA proximity mediated by long, almost perfect stretches of complementary nucleotides rather than RBPs (*15*). The pattern we observe in *PHF20L1* and *CASK* genes and in some other genes with known RNA structure such as SF1 (*41*) suggests that functional PCCRs are supported by RNA contacts from the outer and from the inner side (Figures S7A and S8A). However, we inevitably had to aggregate different biological conditions together because RIC-seq contacts are highly sparse and variable (Figures S3 and S1). Thus, our predictions focus on the core RNA structure formed by PCCRs that is shared by all cell lines. At the same time, we also present evidence of correlated changes between RNA structure and alternative splicing suggesting that the same RNA fragment may be folded and, consequently, spliced differently in different cellular contexts (Figure 3B).

The relationship between intramolecular RNA contacts and complementarity manifests itself in different PCCR features such as higher stability, significance of compensatory substitutions, high abundance of A-to-I editing sites or forked eCLIP peaks. All these features are hallmarks of functional RNA structures, which correspond to PCCRs with substantially lower false discovery rate (*15*). Furthermore, the RF classifier based on RNA contacts confirmed that the most important features are located nearby PCCRs. However, since the number of true positive functional RNA structures in human genes is quite limited, currently there is not sufficient data to estimate the error rate of this approach. For instance, we also attempted to identify RIC-seq contacts in structured RNA classes such as miRNA precursors, however visual inspection revealed that most of them possess only support for the inner contact corresponding to the apical loop of the hairpin but no outer support as shown in Figure 1B (data not shown).

In regard to identification of RNA structures that impact pre-mRNA splicing, in our analysis there was no *a priori* connection to splicing regulation except the requirement that PCCR must loop out a cassette exon. That is, in selecting long-range RNA structures that are potentially associated with splicing, we had no evidence for their impact on splicing other than exon loop-out, i.e., PCCRs that regulate alternative splicing in *PHF20L1* and *CASK* genes were identified by serendipity. We probed many other PCCRs with RIC-seq support as high as that in *PHF20L1* and *CASK*, however without apparent impact on splicing (data not shown). The strategy based on AONs to first probe cis-regulatory signals and, in the case of response, to use minigene mutagenesis appears to be the best at the current state of the art.

The functional relevance of the exon skipping events in *PHF20L1* and *CASK* and their possible regulation deserve further investigation. Here we can only speculate that exon 6 skipping in *PHF20L1* could be related to modulating the function of its TUDOR domain that is encoded within exons 4 and 5. In *CASK*, the inclusion of exon 19 (69 bp) or exon 20 (36 bp) in murine neurons can be induced by KCl treatment, which mimics neuronal excitation (*37*). It was hypothesized that different splice isoforms, including exon 19 skipping isoform, have altered binding properties to *CASK* partners during development stages, as well as in different cell populations with distinct neuronal activity (*38*). These examples demonstrate the importance of the resource provided in this work, which has many practical applications to uncovering new regulatory roles of RNA structures in human disease (*7*).

## Conclusion

We characterized pairs of conserved complementary regions (PCCRs) in terms of support by nearby RNA contacts inferred from RIC-seq experiments in eight human cell lines. Based on this characterization, we identified long-range RNA structures in the human *PHF20L1* gene and in the human and murine *CASK* gene that regulate exon skipping. We provide the results of this work in the form of supplementary data files listing RIC-seq support for PCCRs and a Genome Browser track hub for visualization of the respective RNA contacts.

## Methods

### RIC-seq and RNA-seq experiments

The results of RIC-seq experiments in eight human cell lines including GM12878, H1, HeLa, HepG2, IMR90, K562, hNPC, and neuronal cells (two bioreplicates each) were downloaded from the Gene Expression Omnibus under the accession number GSE190214 in FASTQ format. The matched control RNA-seq experiments were downloaded from the ENCODE consortium under the accession numbers listed in Table S3. RIC-seq and the matched RNA-seq data were processed by RNAcontacts pipeline (*32*) using February 2009 (hg19) assembly of the human genome and GENCODE transcript annotation v41lift37, which were downloaded from Genome Reference Consortium (*42*) and GENCODE website (*43*), respectively. The pipeline generated a track hub for UCSC Genome Browser (*44, 45*) that is available through the link https://github.com/smargasyuk/PHRIC-hub.

### Splicing quantification and analysis

RNA-seq experiments were processed by IPSA pipeline to obtain split read counts supporting splice junctions (*46*). Split read counts were processed with the default settings including entropy content, annotation status and canonical GT/AG dinucleotides at splice sites. The exon inclusion rate (Ψ, PSI, or Percent-Spliced-In) was calculated according to the equation

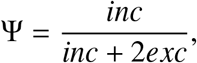

where *inc* is the number of reads supporting exon inclusion and *exc* is the number of reads supporting exon exclusion. Ψ values with the denominator below 10 were considered unreliable and discarded.

Each PCCR containing at least one exon was characterized by the average Ψ value of all exons inside it in each cell line *i*, denoted by Ψ_*i*_. PCCRs with alternatively spliced exons were defined as those having at least two distinct Ψ_*i*_ values for different *i*. We computed the average Ψ_*i*_ across all *i* to characterize the general tendency for exon skipping as a function of the number of cell lines with RIC-seq support. Similarly, we selected PCCR that are supported by at least four reads in at least three cell lines, in which Ψ_*i*_ is also defined, and computed Ψ_*h*_, the average Ψ_*i*_ across cell lines with RIC-seq support, Ψ_*l*_, the average Ψ_*i*_ across cell lines without RIC-seq support, and their difference ΔΨ = Ψ_*h*_ − Ψ_*l*_.

To construct a statistical model, we computed the number of RIC-seq split reads supporting contacts between 50-nt bins −3*L*, −2*L*, −1*L*, 1*L*, 2*L*, 3*L* and −3*R*, −2*R*, −1*R*, 1*R*, 2*R*, 3*R* for each PCCR (Figure 4A). The model was implemented using `RandomForestClassifier` routine from `scikit-learn` Python package v1.1.1 using all PCCRs with at least one contact. The response variable (forked eCLIP peak) was modeled as a function of the number of reads supporting each pair of bins and the spread of the PCCR. At that, the dataset was split into the training set (66%) and the test set (33%) to estimate True Positive Rate (TPR) and False Positive Rate (FPR). The random forest classifier was used at the default settings: 100 trees forest optimizing Gini impurity; tree depths were selected dynamically (leaves are expanded until they are pure or contain only 1 sample).

### Cell culture and transfection

Human A549 lung adenocarcinoma cells and NIH 3T3 mouse fibroblasts were cultured in Dulbecco’s modified Eagle’s medium/Nutrient Mixture F-12 supplemented with 10% fetal bovine serum and 1% GlutaMAX (Thermo Fisher Scientific). 1.5 · 10^5^ cells were seeded in a 12-well plate. One thousand or five hundred nanograms of WT or mutated minigene plasmids were transfected into NIH 3T3 or A549 cells using Lipofectamine 3000 (Invitrogen) for 24 hours. AON (13-mer) transfection was performed with Lipofectamine RNAiMAX (Invitrogen) in OptiMEM serum-reduced media (Gibco) for 48 hours.

### Antisense oligonucleotides

All antisense oligonucleotides (AONs) were designed as locked nucleic acid (LNA)-based with a DNA substitution at every second nucleotide (*47*). Synthesis of LNA/DNA mixmers was carried out by Syntol JSC (Moscow, Russia). Sequences of AONs are listed in Table S4.

### Minigene construction and mutagenesis

Minigenes encoding *CASK* exons 18–20 and *PHF20L1* exons 5–7 were cloned into a pRK5 expression vector containing CMV promoter for the evaluation of target exon inclusion rates. The minigene sequence was amplified from A549 or NIH 3T3 genomic DNA using Q5 High-Fidelity DNA polymerase (New England Biolab). *PHF20L1* minigene was assembled using restriction-free cloning. The human *CASK* fragment of DNA was cloned using blunt end cloning protocol. The murine *CASK* was cloned into the ClaI and SalI sites of the pRK5 vector. All mutations in minigenes were introduced by PCR-based site-directed mutagenesis. All primers for cloning and mutagenesis are listed in Table S5 and Table S6, respectively. All constructs were confirmed by sequencing.

### RNA extraction

Total RNA was isolated by a guanidinium thiocyanate phenol-chloroform method using ExtractRNA Reagent (Evrogene) or PureLink RNA minikit (Invitrogen) (*48*). One thousand ng of total RNA was first subjected to RNase-Free DNase I digestion (Thermo Fisher Scientific) at 37°C for 30 minutes to remove contaminating gDNA. Next, 500 ng of total RNA was used for cDNA synthesis using Maxima First Strand cDNA Synthesis Kit for RT-qPCR (Thermo Fisher Scientific) to a final volume of 10 μl. cDNA was diluted 1:10 with nuclease free water for qPCR and RT-PCR analysis.

### RT-PCR

RT-PCR analysis was used for all human *CASK* experiments to assess the ratio of splice isoforms. Reactions were carried out using PCR Master Mix (2x) (Thermo Scientific). RT-PCR was carried out under the following conditions: 95°C for 3 min, 35 cycles at 95°C for 30s, 54°C for 30 s, 72°C for 1 min, ending at 72°C for 5 min. The resulting products were analyzed on a 3% agarose gel stained with ethidium bromide and visualized using ChemiDoc XRS+ (Bio-Rad). The relative amounts of the resulting products were analyzed by scanning the gels and determining the intensities of ethidium bromide stained bands using Image Lab software Version 1.36b (Bio-Rad). Endpoint PCR for all mouse *CASK* experiments was performed using 0.25 U recombinant Taq DNA Polymerase (Thermo Fisher Scientific) with 2 mM MgCl_2_, 1x KCl buffer, 2μl of cDNA (same as for qPCR), 0.2 mM of each dNTPs, and 420nM of each forward and reverse primer (Table S7). Cycling conditions were as follows: 95°C for 5 min, followed by 35 cycles at 95°C for 30 s, 61°C for 30 s, and 72°C for 30 s and ended at 72°C for 5 min. PCR products were visualized on the agarose gel.

### qPCR

qPCR reactions were performed in triplicates in a final volume of 12μl in 96-well plates with 420 nM gene-specific primers, 2μl of cDNA using Maxima SYBR Green/ROX qPCR Master Mix (2X) (Thermo Fisher Scientific). A sample without reverse transcriptase enzyme was included as control to verify the absence of genomic DNA contamination. Amplification of the targets was carried out on CFX96 Real-Time system (Bio-Rad), cycle parameters were as follows: 95°C for 2 min, followed by 45 cycles at 95°C for 30 s, 61°C for 30 s, and 72°C for 30 s, ending at 72°C for 5 min. The expression of genes and gene isoforms was calculated using the assay specific PCR efficiency. The result of replicate measurement was considered an outlier and rejected if its *C_q_* value was not in the range of 0.5 cycles (*49*).

## Availability of data and materials

The data obtained in this work (Supplementary Data Files 1 and 2) are available through Zenodo repository (*50*).

## Authors’ contributions

DP designed and supervised the study; DP and SM performed data analysis; CC provided RIC-seq data; MK, MP, and DS performed the experiments; DP, SM, and MP wrote the first draft of the manuscript. All authors edited the final version of the manuscript.

## Acknowledgments

The authors thank Margarita Vorobiova for feedback on the manuscript. All authors acknowledge Prof. O.A. Dontsova and Prof. A.A. Mironov for insightful discussions.

